# Fungal dual-domain LysM effectors undergo chitin-induced intermolecular, and not intramolecular, dimerization

**DOI:** 10.1101/2020.06.11.146639

**Authors:** Hui Tian, Gabriel L. Fiorin, Anja Kombrink, Jeroen R. Mesters, Bart P.H.J. Thomma

## Abstract

Chitin is a homopolymer of β-(1,4)-linked *N*-acetyl-D-glucosamine (GlcNAc) and a major structural component of fungal cell walls. In plants, chitin acts as a microbe-associated molecular pattern (MAMP) that is recognized by lysin motif (LysM)-containing plant cell surface-localized pattern recognition receptors (PRRs) that activate a plethora of downstream immune responses. In order to deregulate chitin-induced plant immunity and successfully establish infection, many fungal pathogens secrete LysM domain-containing effector proteins during host colonization. It was previously shown that the LysM effector Ecp6 from the tomato leaf mould fungus *Cladosporium fulvum* can outcompete plant PRRs for chitin binding because two of its three LysM domains cooperate to form a composite groove with ultra-high (pM) chitin-binding affinity. However, most functionally characterized LysM effectors contain only two LysMs, including *Magnaporthe oryzae* MoSlp1, *Verticillium dahliae* Vd2LysM, and *Colletotrichum higginsianum* ChElp1 and ChElp2. Here, we performed modelling, structural and functional analyses to investigate whether such dual-domain LysM effectors can also form ultra-high chitin-binding affinity grooves through intramolecular LysM dimerization. However, our study suggests that intramolecular LysM dimerization does not occur. Rather, our data support the occurrence of intermolecular LysM dimerization for these effectors, associated with a significantly lower chitin binding affinity than monitored for Ecp6. Interestingly, the intermolecular LysM dimerization allows for the formation of polymeric complexes in the presence of chitin. Possibly, such polymers may precipitate at infection sites in order to eliminate chitin oligomers, and thus suppress the activation of chitin-induced plant immunity.

## INTRODUCTION

Chitin is a homopolymer of β-(1,4)-linked *N*-acetyl-D-glucosamine (GlcNAc) and a major structural component of fungal cell walls (Lenardon et al., 2010; Free, 2013). In plants, chitin has been characterized as a fungal microbe-associated molecular pattern (MAMP) that can be recognized by cell surface-localized pattern recognition receptors that contain extracellular lysin motif (LysM) domains (LysM-PRRs) (Zhang et al., 2007; Zipfel, 2008; Sanchez-Vallet et al., 2015; Rovenich et al., 2016). Upon recognition of chitin by such receptors, plants evoke a broad range of immune responses, including the generation of ion fluxes, the production of reactive oxygen species (ROS), the activation of mitogen-associated protein kinases (MAPKs) and the expression of defence-related genes that include those encoding hydrolytic enzymes, such as chitinases, in order to halt fungal invasion (Felix et al., 1993; Jones and Dangl, 2006; Altenbach and Robatzek, 2007; Boller and Felix, 2009; Sanchez-Vallet et al., 2015). LysM-PRRs have been functionally characterized in several plants, including the model plant Arabidopsis (*Arabidopsis thaliana*) in which the LysM receptor AtLYK5 was reported to bind chitin with high affinity (1.72 µM) and form a tripartite receptor complex with AtLYK4 and AtCERK1 to initiate chitin signalling (Cao et al., 2014). AtCERK1 was found to bind chitin directly as well, albeit with significantly lower affinity than AtLYK5 (Miya et al., 2007; Petutschnig et al., 2010; Cao et al., 2014).

To overcome chitin-induced immunity, successful fungal pathogens evolved various strategies to either protect fungal cell wall chitin against hydrolysis by host enzymes, or prevent the activation of plant immunity by fungal cell wall-derived chitin oligomers (de Jonge et al., 2010; Rovenich et al., 2014; Sanchez-Vallet et al., 2015). For the fungus *Cladosporium fulvum*, causal agent of leaf mould disease of tomato, several strategies to deal with chitin-triggered immunity have been characterized. During host colonization, *C. fulvum* secretes the effector protein Ecp6 that contains three LysMs and binds chitin oligosaccharides with ultra-high affinity, leading to the suppression of chitin-induced plant immune responses (Bolton et al., 2008; de Jonge et al., 2010). A crystal structure of Ecp6 revealed that two of its three LysMs cooperate to form a composite chitin-binding groove that binds chitin through intrachain LysM dimerization (Sanchez-Vallet et al., 2013). *Zymoseptoria tritici*, the causal agent of Septoria tritici blotch (STB) of wheat, secretes the LysM effector Mg3LysM, a close homolog of Ecp6, that similarly suppresses chitin-triggered immunity (Marshall et al., 2011). Additionally, *Z. tritici* secretes two effectors that comprise a single LysM only, Mg1LysM and Mgx1LysM that were both characterized to protect hyphae against hydrolysis by plant chitinases and to suppress chitin-triggered immunity in plants (Marshall et al., 2011; Tian et al., 2021). An Mg1LysM crystal structure showed that two Mg1LysM monomers form a chitin-independent homodimer via the β-sheet in the *N*-terminus of Mg1LysM (Sánchez-Vallet et al., 2019). Furthermore, Mg1LysM homodimers were shown to undergo ligand-induced polymerization in the presence of chitin, leading to a polymeric structure that is able to protect fungal cell walls (Sánchez-Vallet et al., 2019). Similar chitin-induced homopolymer formation was shown for Mgx1LysM (Tian et al., 2021).

Suppression of chitin-triggered immunity by secreted fungal effectors that carry no other recognizable domains than LysM domains, collectively referred to as LysM effectors, has been demonstrated for various phytopathogenic fungi. For instance, *Magnaporthe oryzae*, the causal agent of rice blast disease, secretes the LysM effector Slp1 to bind chitin and suppress chitin-triggered immune responses (Mentlak et al., 2012). Similarly, the Brassicaceae anthracnose fungus *Colletotrichum higginsianum* secretes ChElp1 and ChElp2, while the broad host-range vascular wilt fungus *Verticillium dahliae* secretes Vd2LysM (Takahara et al., 2016; Kombrink et al., 2017). While these examples are from plant-associated Ascomycete fungi, also plant-associated fungi that belong to other phyla utilize LysM effectors to suppress chitin-triggered immunity. For instance, the Basidiomycota soil-borne broad host-range pathogen *Rhizoctonia solani* secretes RsLysM, while the Glomeromycota arbuscular mycorrhizal fungus *Rhizophagus irregularis* secretes RiSLM to suppress chitin-triggered immunity (Dolfors et al., 2019; Zeng et al., 2020). The latter example demonstrates that also non-pathogenic fungi utilize LysM effectors in their interactions with host plants. Moreover, the finding that LysM effectors contribute to the virulence of the Ascomycete fungus *Beauveria bassiana* by evasion of immune responses in insect hosts demonstrates that LysM effectors play roles in fungal interactions beyond plant hosts (Kombrink and Thomma, 2013; Cen et al., 2017).

Based on the functional analysis of *C. fulvum* Ecp6, it has been proposed that the ability to suppress chitin-triggered immunity resides in the ability to form a composite chitin-binding groove through intrachain LysM dimerization to bind chitin with ultrahigh affinity, such that host chitin receptors are outcompeted for substrate binding (Sanchez-Vallet et al., 2013; Sanchez-Vallet et al., 2015). However, most LysM effectors that are functionally characterized today contain only two LysMs; so-called dual-domain LysM effectors. It has remained unclear thus far whether these dual-domain LysM effectors are able to undergo intramolecular LysM dimerization, like in Ecp6, to establish ultrahigh affinity chitin-binding sites, or whether chitin binding by these effectors is based on another mechanism.

## RESULTS

### Structural prediction precludes intrachain LysM dimerization

It has previously been determined that *M. oryzae* MoSlp1, *V. dahliae* Vd2LysM, and *C. higginsianum* ChElp1 and ChElp2 comprise two LysM domains, bind chitin and suppress chitin-induced host immunity (Mentlak et al., 2012; Takahara et al., 2016; Kombrink et al., 2017). Their length ranges from 145 aa (for Vd2LysM) to 176 aa (for ChElp2), with the molecular weight of the mature proteins ranging from 14.24 to 16.14 kDa (Fig. 1A). An amino acid sequence alignment of their LysM domains with those of *C. fulvum* Ecp6 displayed significant conservation of the domains, and of the residues involved in chitin binding in particular (Fig. 1B). LysM domains are compact modules that display a β-α-α-β secondary structure motif (Bateman and Bycroft, 2000) in which the termini of the beta sheets are typically kept together by a disulfide bond at the LysM domain borders. In Ecp6, three disulfide bonds are found within the individual LysM domains (LysM1: C7-C61; LysM2: C80-C134; LysM3: C139-C191) and a fourth bond (C35-C69) locates in between LysM1 and the linker between the two LysMs (Fig. 1C). Accordingly, in MoSlp1, Vd2LysM, ChElp1 and ChElp2 the cysteines to form a disulfide bond within each of the individual LysM domains are conserved and, similar to Ecp6, those to generate another disulfide bond in between LysM1 and the linker between the two LysM domains (MoSlp1: C46-C80; Vd2LysM: C31-C69; ChElp1: C56-C90; ChElp2: C62-C96) are conserved as well (Fig. 1C). Intriguingly, these latter disulfide bonds will anchor a significant part of the linker between the two LysM domains to the beginning of the second α-helix of LysM1 in each of these effectors, severely limiting the part of the linker that remains free to rotate. This is the most extreme for Vd2LysM, for which ten of the 19 amino acids of the linker between the two LysMs become fixed and a flexible stretch of only nine amino acids remains (Fig. 1C).

**Figure 1.**
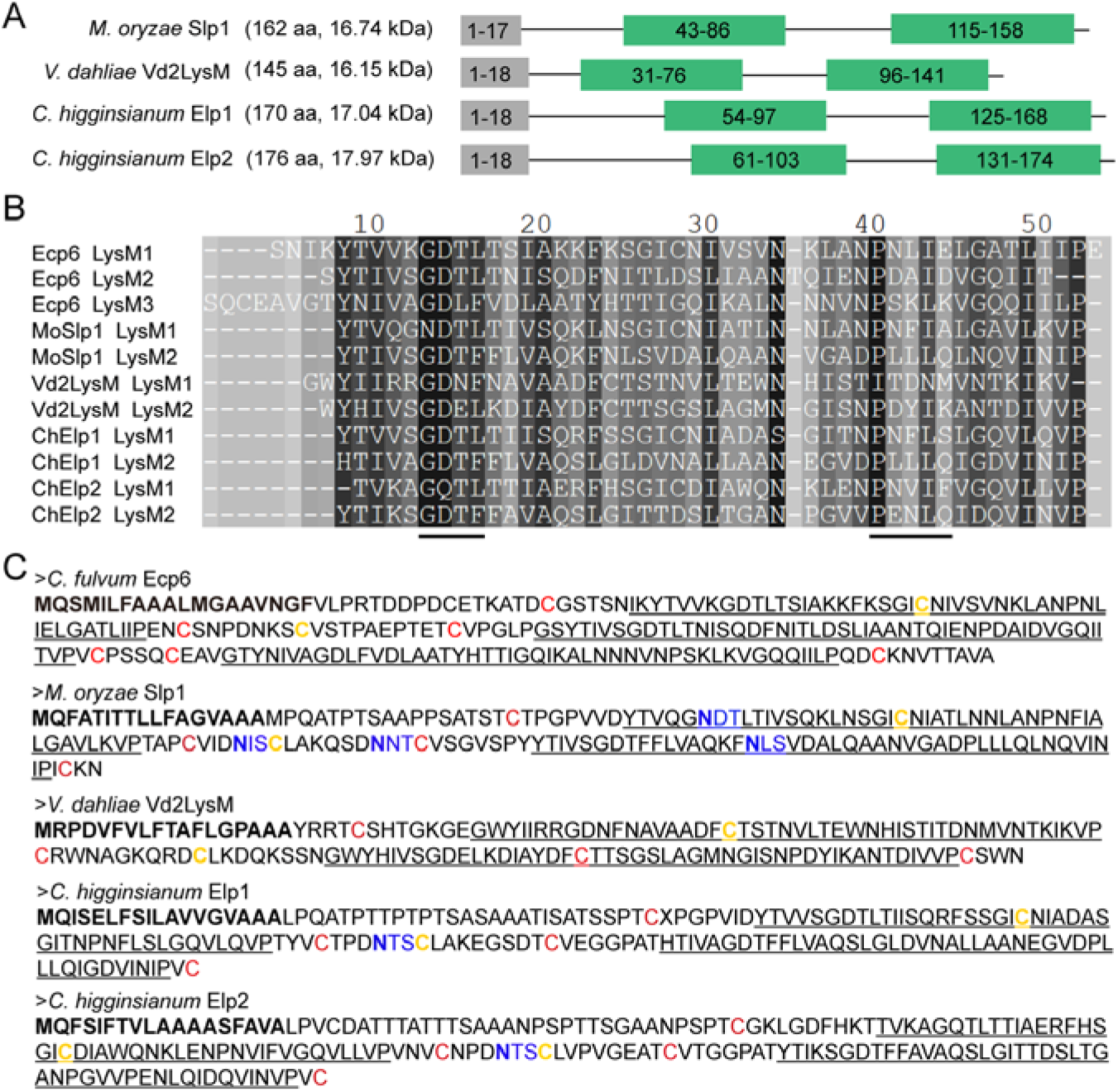
Conservation of chitin binding sites and of disulfide bond formation among Ecp6 and the four dual-domain LysM effectors. (A) Schematic representation of the four fungal dual-domain LysM effectors: *Magnaporthe oryzae* MoSlp1, *Verticillium dahliae* Vd2LysM, and *Colletotrichum higginsianum* ChElp1 and ChElp2. Signal peptides (grey boxes) were predicted with SignalP 4.0 (http://www.cbs.dtu.dk/services/SignalP-4.0/) and LysM domains (green boxes) with InterPro (https://www.ebi.ac.uk/interpro/). The amino acid numbers that compose the motif are indicated in the boxes. (B) Amino acid sequence alignment of the LysM domains of MoSlp1, Vd2LysM, ChElp1 and ChElp2 in comparison with those of Ecp6. The black lines indicate chitin-binding sites. (C) Amino acid sequences of Ecp6, MoSlp1, Vd2LysM, ChElp1 and ChElp2 with signal peptides in bold, LysM domains underlined and the putative N-glycosylation sites highlighted in blue with the asparagines that may be N-glycosylated in blue bold font. Cysteine residues involved in disulfide bond formation are indicated red and green with those involved in disulfide bonds within individual LysM domains in red, and those involved in the bond between LysM1 and the linkers between the two LysM domains in yellow.

To assess whether intramolecular LysM dimerization can occur in the dual-domain LysM effectors, all four proteins were modelled based on the LysM1-LysM3 composite chitin binding groove of Ecp6 using Phyre2, I-TASSER and AlphaFold (Roy et al., 2010; Yang and Zhang, 2015; Kelley et al., 2015; Jumper et al., 2021). Phyre2 successfully superposed the LysM domains of each of the effectors on LysM1 and LysM3 of Ecp6. Moreover, the software successfully predicted the disulfide bond that anchors a significant part of the linker between the two LysM domains to the beginning of the second α-helix of LysM1 for all LysM effectors but Vd2LysM (Fig. 2). However, all four structure predictions displayed a disruption of the linker between the LysM domains (Fig. 2), suggesting that (the flexible part of) the linkers of MoSlp1, Vd2LysM, ChElp1 and ChElp2 are too short to accommodate intramolecular LysM dimerization. Accordingly, an artificial insertion of a 5-amino acid sequence (GGSGG) in the linker of ChElp2 led to predicted intramolecular LysM dimerization in a structure model with an uninterrupted loop and a root-mean-square deviation (r.m.s.d.) value of 0.16 Å for 499 atom pairs (Fig. 2).

**Figure 2.**
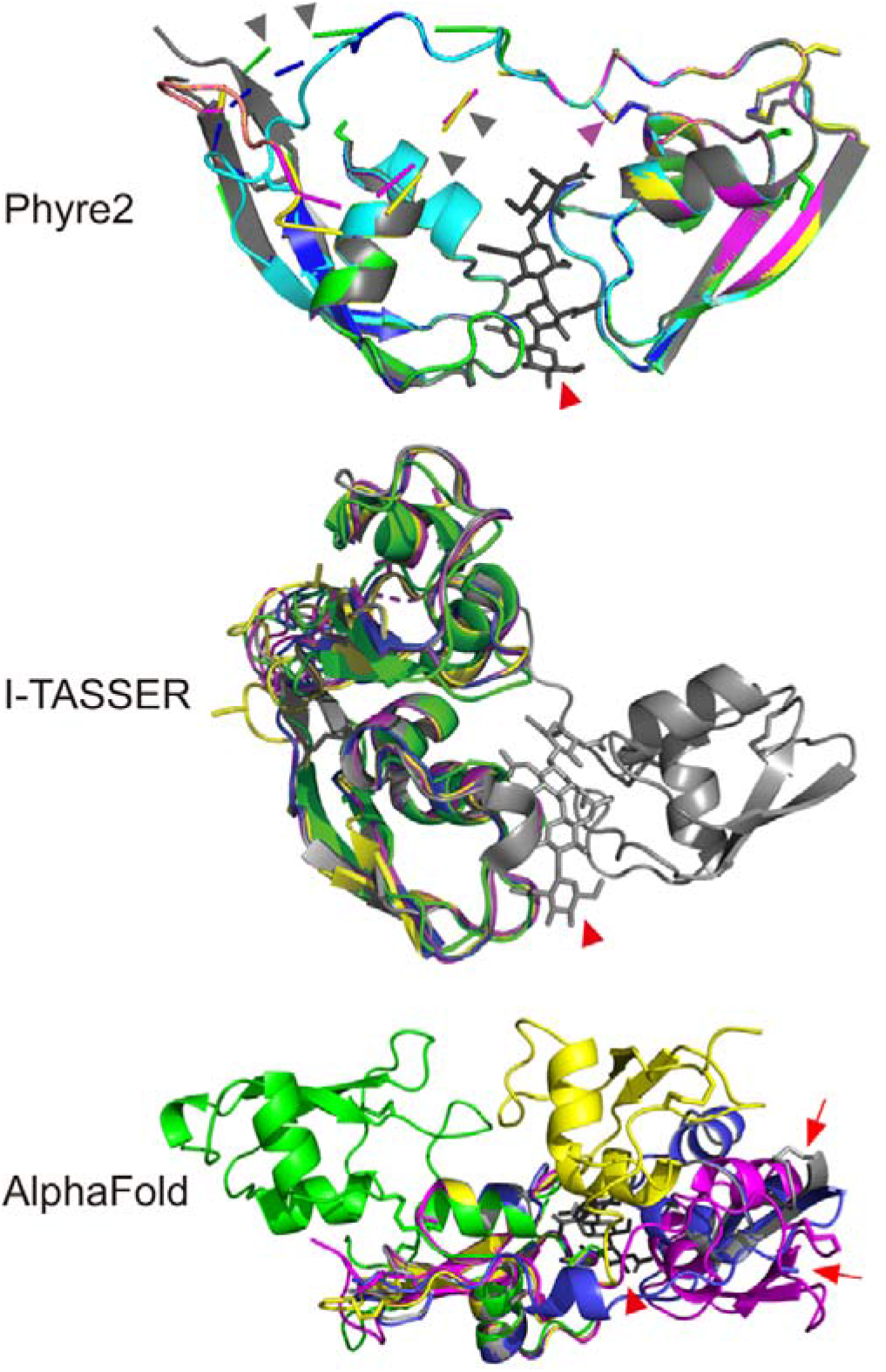
*In-silico* prediction of the three-dimensional structures of MoSlp1, Vd2LysM, ChElp1 and ChElp2. The prediction was conducted based on the composite chitin binding groove composed of LysM1 and LysM3 of Ecp6 (Sánchez-Vallet et al., 2013) with Ecp6 in grey, MoSlp1 in yellow, Vd2LysM in green, ChElp1 in blue, and ChElp2 in purple. Furthermore, for the Phyre2 prediction, ChElp2 carrying an artificial insertion of 5-amino acid (GGSGG) in the linker between the two LysMs is indicated in cyan. Red arrowheads indicate a chitin tetramer in the Ecp6 LysM1-LysM3 binding groove. Purple and grey arrowheads indicate predicted disulphide bonds for MoSlp1, ChElp1 and ChElp2 and disrupted linkers for the four dual-domain LysM effectors, respectively. The red arrows indicate the questionable offset position of C99 in the ChElp1 AlphaFold predicted structure (blue) and of C139 in the crystal structure of Ecp6 (grey). All structures were visualized using the PyMOL molecular graphics system (Schrodinger LLC, 2015).

The I-TASSER prediction did not result in the display of disrupted linkers. However, whereas the modelling was attempted on the composite binding groove of Ecp6 composed by LysM1 and LysM3, I-TASSER modelled the structure of the LysM effectors on LysM2 and LysM3 of Ecp6 (Fig. 2), r.m.s.d. values ranging from 0.49 Å (442 atom pairs) for ChElp2, 0.60 Å (470 atom pairs) for ChElp21, 0.64 Å (508 atom pairs) for MoSlp1 to 1.67 Å (395 atom pairs) for Vd2LysM. Thus, like Phyre2, also the I-TASSER modeling suggests that intramolecular LysM dimerization is not likely.

Also the third prediction software, AlphaFold, yielded structure models that were not compatible with intramolecular chitin binding, with r.m.s.d. values as high as 2.62 Å (567 atom pairs) and 5.42 Å (602 atom pairs) for ChElp2 and MoSlp1, respectively (Fig. 2). The ChElp1 structure model appeared compatible with intramolecular chitin binding displaying an overall r.m.s.d. value of 0.153 Å (541 atom pairs). However, the low r.m.s.d. value was only attainable because none of the anticipated disulfide bonds (C28-C83, C56-C90 and C99-C152) materialized in the model. Especially C99 appears completely out of position because the last three C-terminal amino acids including C152 are lacking in the model (Fig. 2). The overall lack of cystins in the model permitted the flexible loop between the two LysM domains to take a shortcut route, thereby ending in a dubious model abandoning the basic features of a LysM domain. Surprisingly, in the Vd2LysM model a different cysteine pair occurred, C5-C59, C31-C101 and C69-C124, resulting in a model in which the two chitin binding motifs are pointing in opposite directions. Taken together, all predictions based on the various software packages do not support intramolecular chitin binding.

### Crystallization attempts for the LysM effectors failed

The most direct method to reveal the chitin-binding mechanism of a LysM effector is by determination of a three-dimensional protein structure in the presence of chitin, for instance by X-ray crystallography, as previously performed for the single-domain LysM effector Mg1LysM (Sánchez-Vallet et al., 2020) and the three-domain LysM effector Ecp6 (Sánchez-Vallet et al., 2013). Thus, we pursued this strategy for the dual-domain LysM effectors as well. This strategy requires a protein crystal of sufficient size and quality to be used in an X-ray diffraction experiment, which in turn requires highly pure protein of a sufficiently high concentration. To this end, the four dual-domain LysM effectors were heterologously produced in the yeast *Pichia pastoris* (Fig. S1A). Whereas MoSlp1 possesses four potential glycosylation sites (**N**^48^DT, **N**^94^IS, **N**^130^LS and **N**^104^NT), ChElp1 and ChElp2 contain only a single potential glycosylation site (**N**^105^TS and **N**^111^TS) and Vd2LysM is not predicted to possess any glycosylation site (Fig. 1C) which corresponds to findings in a glycoprotein staining assay (Fig. S1B). Nevertheless, attempts to remove potential glycan chains did not lead to decreased molecular sizes of MoSlp1, Vd2LysM and ChElp2 while fetuin as positive control revealed a clear size shift already at 2 h after incubation (Fig. S1C). As we have previously successfully crystallized Ecp6 protein that was produced in the same manner despite containing two spatially close glycosylation sites that were indeed found to be glycosylated in the crystal structure (Sanchez-Vallet et al., 2013), we decided to continue with these protein preparations. Also, the molecular homogeneity of the protein solutions was improved by gel filtration combined with the treatment with the nonionic detergent decyl β-D-maltopyranoside (DM) (Fig. S2). Subsequently, all four proteins were subjected to extensive crystallization screenings. In addition, ChElp2 and Vd2LysM were produced in *Escherichia coli* and subjected to crystallization screenings as well. Unfortunately, none of the conditions tested resulted in a genuine protein crystal for any of the dual-domain LysM effectors. Although crystallization is an empirical process and there are many reasons why no crystals are obtained, we reasoned that the lack of intramolecular chitin binding may also hamper crystal growth.

### Chitin-induced intermolecular dimerization leads to LysM effector polymerization

As all our crystallization attempts for the four dual-domain LysM effectors failed, we pursued other strategies to provide evidence for the mechanism of chitin binding. We reasoned that treatment with chitin oligomers would lead to higher order oligomeric or polymeric protein complexes if intermolecular LysM dimerization occurs (Fig. 3A, hypothesis I), while such complexes will not be formed in case of intramolecular LysM dimerization (Fig. 3A, hypothesis II). To address these alternative hypotheses, ChElp2 was selected as a representative and expressed in *E. coli* to obtain protein that is devoid of chitin. ChElp2 was produced either with a His-sumo-tag (14 kDa) or with a FLAG-tag (1 kDa) and both forms were incubated together in the absence and presence of chitin. His-sumo-tagged ChElp2 was pulled-down with anti-His magnetic beads and the subsequent western blot (WB) result showed that FLAG-tagged ChElp2 was associated with His-sumo-tagged ChElp2 only in the presence of chitin (Fig. 3B), indicating chitin-induced intermolecular interactions between ChElp2 monomers.

**Figure 3.**
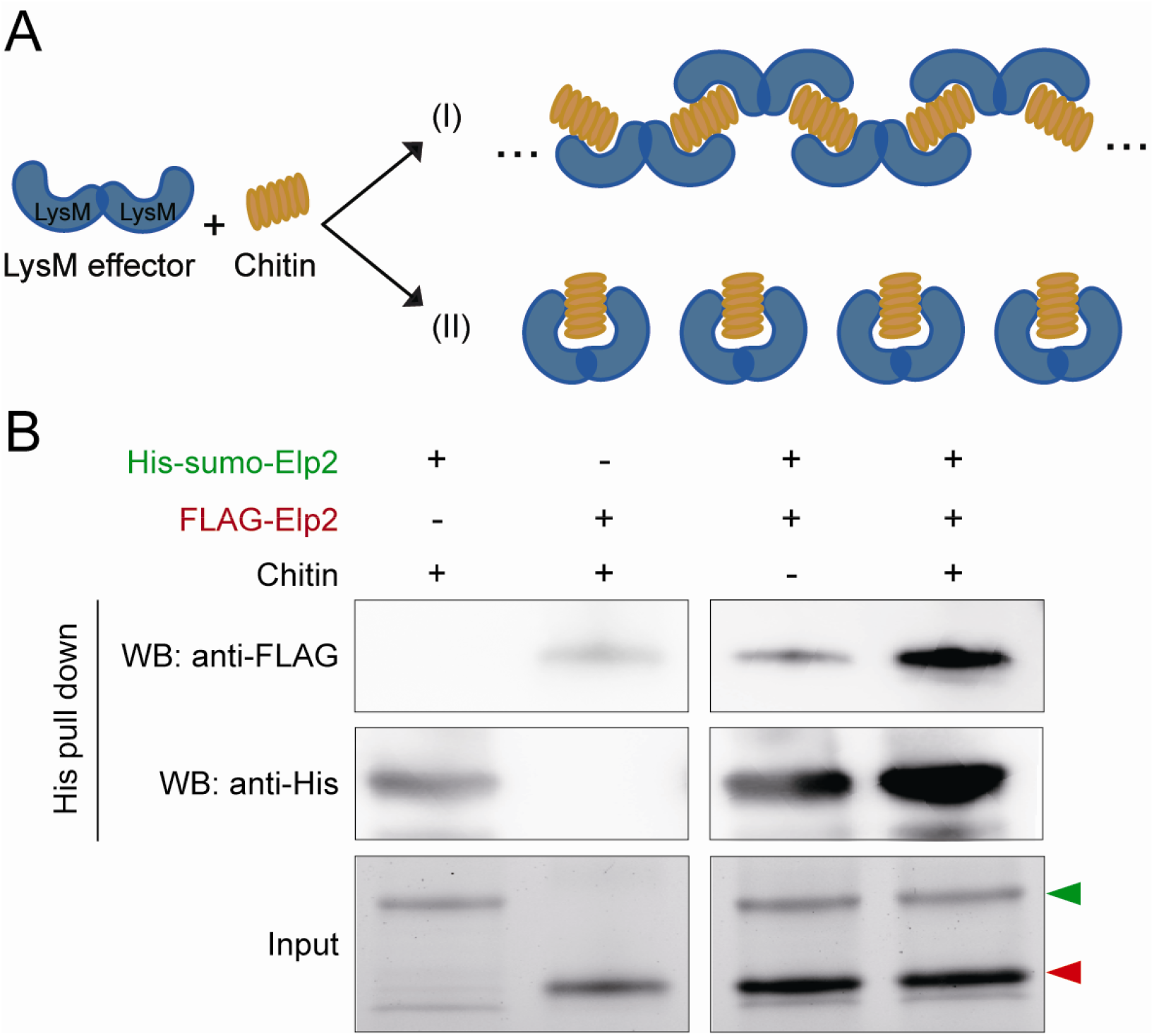
ChElp2 binds chitin via intermolecular LysM dimerization. (A) Two hypotheses for chitin binding by dual-domain LysM effectors that may bind chitin through: (I) intermolecular dimerization in which LysM effectors may undergo ligand-induced polymerization, or through: (II) intramolecular dimerization that should not lead to polymerization. (B) Escherichia coli-produced His-sumo-ChElp2 and FLAG-ChElp2 were incubated either individually in the presence of chitin or together in the absence and presence of chitin for 4-6 hours. His-sumo-ChElp2 was pulled-down by anti-His magnetic beads and the subsequent western blot analysis was performed with either anti-FLAG or anti-His antibody. Green and red arrow heads indicate the molecular sizes of His-sumo-Elp2 and FLAG-Elp2 on protein polyacrylamide gel, respectively.

To provide further evidence for chitin-induced intermolecular interactions, we assessed the ChElp2 particle size distribution in solution upon chitin addition with dynamic light scattering (DLS). Chitin addition at a molar ratio of 1:3 (protein:chitin) shifted a portion of ChElp2 particles from around 10 nm in the absence of chitin to 100 nm in the presence of chitin (Fig. 4A). Further increase of exogenously added chitin to a molar ratio of 1:6 significantly reduced the signal at 10 nm while inducing a strong signal at 10 μm (Fig. 4A), indicative of chitin-induced polymerization of ChElp2. To demonstrate the requirement of chitin binding for the observed polymerization, we produced a ChElp2 mutant in which the threonine within each of the LysM domains that is critical for chitin binding (LysM1, T49; LysM2, T120) was replaced by tryptophan, produced the mutant in *E. coli*, and confirmed the compromised chitin binding affinity by isothermal titration calorimetry (ITC) (Fig. 4B). Subsequently, the ChElp2^T49W-T120W^ mutant was subjected to DLS analysis in the presence of exogenously added chitin, revealing that the particle size shift that is observed for wild-type ChElp2 was not observed for the mutant (Fig. 4A). Thus, we conclude that chitin binding is required for ChElp2 polymerization.

**Figure 4.**
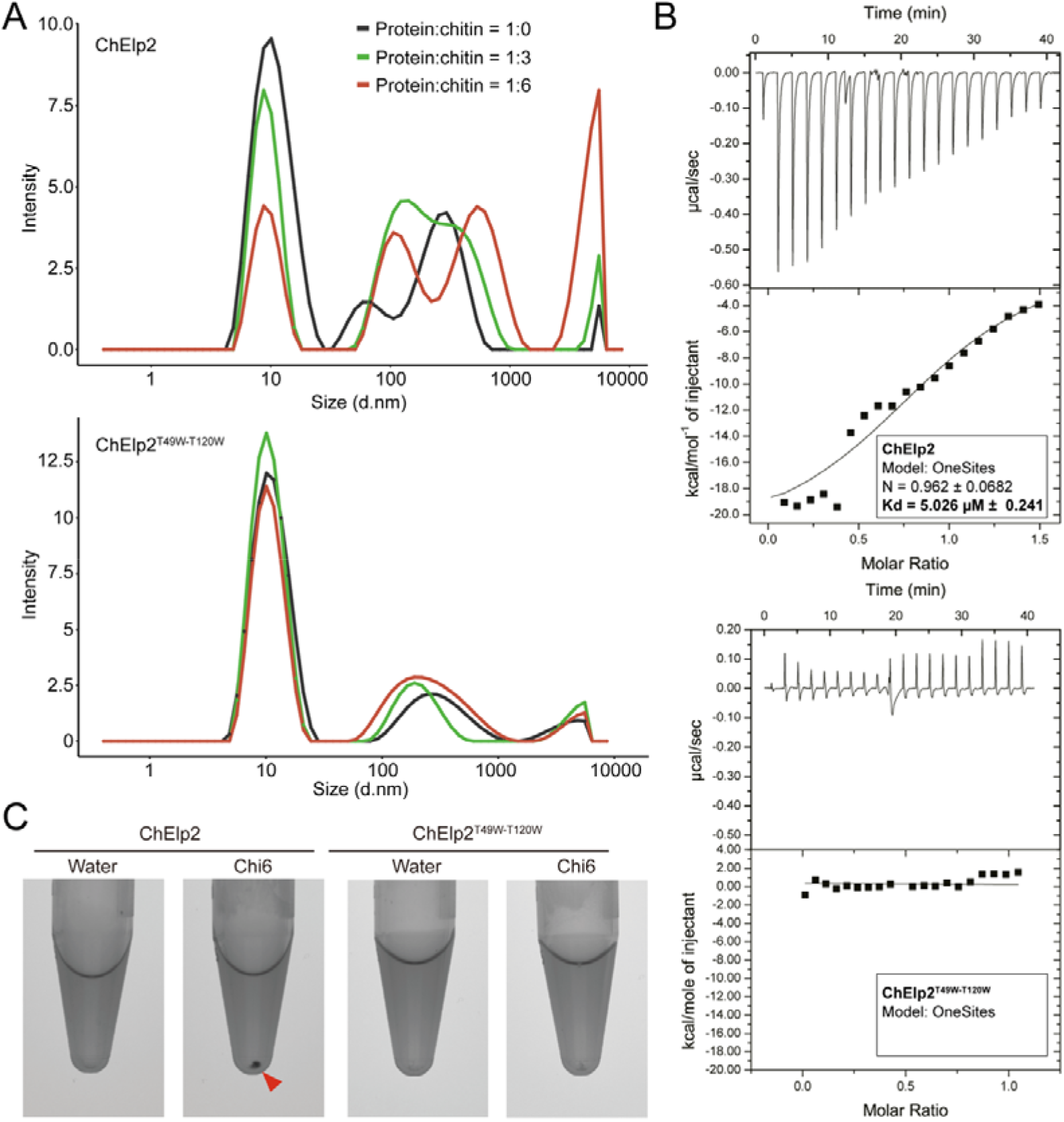
ChElp2 undergoes chitin-induced polymerization. (A) ChElp2 particle size distribution in protein solution with diameter values in nanometers (d.nm) measured by dynamic light scattering (DLS). (B) Raw data and integrated heat measurements from isothermal titration calorimetry of chitohexose binding to wild-type ChElp2 and the mutant ChElp2^T49W-T120W^ in which the threonine within each of the LysM domains that is critical for chitin binding (LysM1, T49; LysM2, T120) was replaced by tryptophan. Lines in the lower panel represent best-fit curves for a one binding-site model. (C) ChElp2 and ChElp2^T49W-T120W^ were incubated with chitin hexamer or water as control. After overnight incubation, methylene blue was added and protein solutions were centrifuged, resulting in protein pellets (red arrowheads) as a consequence of polymerization.

As an additional line of evidence, we hypothesized that if ChElp2 undergoes chitin-induced polymerization, we should be able to precipitate polymeric complexes during centrifugation. Thus, we incubated ChElp2, and ChElp2^T49W-T120W^ and Ecp6 as negative controls, overnight with chitohexaose and subsequently centrifuged the samples at 20,000 g in the presence of 0.002% methylene blue to visualize the protein. As expected, a clear protein pellet appeared when ChElp2 was incubated with chitin, but not in the control treatment without chitin, nor in the ChElp2^T49W-T120W^ and Ecp6 samples (Fig. 4C; Fig. 6). Thus, we confirmed that chitin binding is required for ChElp2 polymerization.

To provide further evidence that specific chitin binding, rather than presence of chitin or other carbohydrates, is required for ChElp2 polymerization, 100 μM ChElp2 was incubated with 1 mM diacetyl-chitobiose (Chi2), triacetyl-chitotriose (Chi3), tetraacetyl-chitotetraose (Chi4), pentaacetyl-chitopentaose (Chi5), hexaacetyl-chitohexaose (Chi6) or glucose. After centrifugation, protein pellets were observed for ChElp2 incubations with Chi3, Chi4, Chi5 and Chi6, but not with Chi2 or glucose (Fig. 5A). Arguably, given their differences in length, equimolar amounts of the various chitin oligomers result in different final amounts of GlcNAc units. Thus, we repeated the assay upon calibrating the various chitin oligomers such that the final amount of GlcNAc units is the same in the various incubations. Also under these conditions, no protein pellets formed upon incubation with Chi2 (Fig. 5B). Considering that a LysM domain binds chitin oligomers that are composed of minimum three GlcNAc units, our findings support that chitin binding, rather than mere chitin addition, is required for pellet formation.

**Figure 5.**
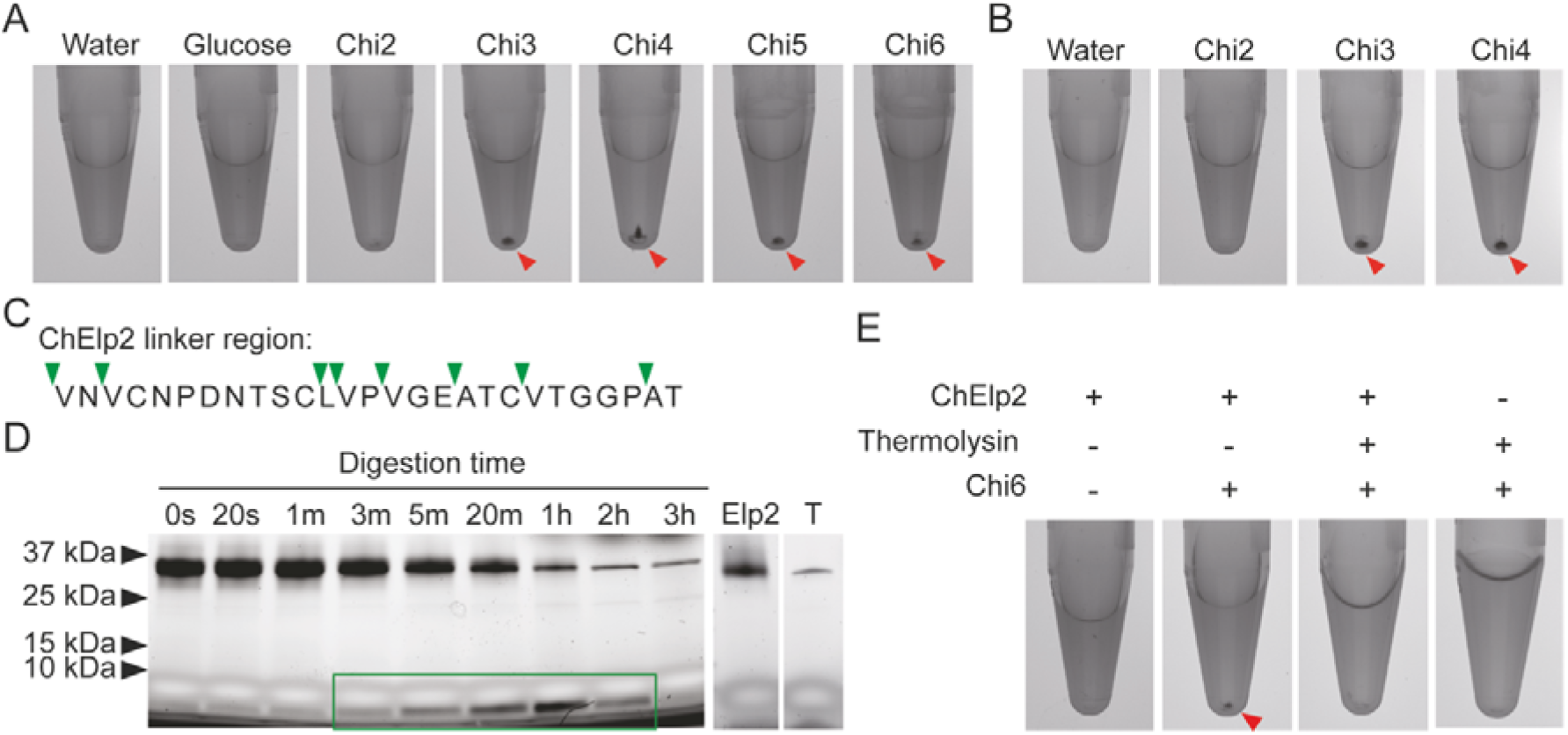
Chitin-binding and the physical association of the two LysM domains are required for ChElp2 polymerization. (A) 100 µM ChElp2 was incubated with 1 mM diacetyl-chitobiose (Chi2), triacetyl-chitotriose (Chi3), tetraacetyl-chitotetraose (Chi4), pentaacetyl-chitopentaose (Chi5), hexaacetyl-chitohexaose (Chi6) or glucose in a total volume of 400 µL. After overnight incubation, methylene blue was added and protein solutions were centrifuged, resulting in protein pellets that are indicated by red arrowheads. (B) 100 µM ChElp2 was incubated with Chi2 (3 mM), Chi3 (2 mM) and Chi4 (1.5 mM) in a total volume of 400 µL such that the final amount of GlcNAc units is the same for all incubations. After overnight incubation, methylene blue was added and protein solutions were centrifuged, resulting in protein pellets that are indicated by red arrowheads. (C) Amino acid sequence of the ChElp2 linker between the two LysMs with putative thermolysin cleavage sites indicated by green arrowheads. (D) P. pastoris-produced ChElp2 was incubated with thermolysin (T) for 3 hours, protein samples were collected at different time points and analysed on protein polyacrylamide gel. (E) P. pastoris-produced ChElp2 incubated with or without thermolysin for 1 hour, followed by overnight incubation with chitin hexamer or water. Methylene blue was added and the protein solutions were centrifuged, resulting in protein pellets that are indicated by red arrowheads.

As a final experiment to substantiate the link between chitin binding, intermolecular dimerization and polymerization, we performed limited ChELP2 digestion with thermolysin. Arguably, thermolysin will first of all target the unstructured linker between the LysM domains rather than the highly structured LysM domains themselves (Fig. 5C). Accordingly, after 5 min of thermolysin treatment a single protein band that represents a single LysM domain could already be observed on polyacrylamide gel, which increased in intensity at expense of the band that represents the full-length LysM effector protein up to 1 h after thermolysin addition (Fig. 5D). While a clear protein pellet was observed in untreated ChElp2 upon chitin addition followed by centrifugation, no pellet formed for ChElp2 treated with thermolysin for 1 hr and incubated with chitin (Fig. 5E). Thus, we conclude that the physical association of the LysM domains is required to permit polymerization based on chitin-induced intermolecular LysM dimerization of ChElp2.

To assess whether our conclusions for ChElp2 can be generalized and applies for other dual-domain LysM effectors as well, the polymerization assay in the presence of chitin was performed for *P. pastoris*-produced ChElp2, and for MoSlp1 and Vd2LysM as well. Indeed, clear pellets were formed upon incubation of these proteins with chitin, followed by centrifugation (Fig. 6). Collectively, our findings argue that dual-domain LysM effectors undergo inter-rather than intramolecular dimerization upon chitin binding, leading to polymerization of these LysM effectors.

**Figure 6.**
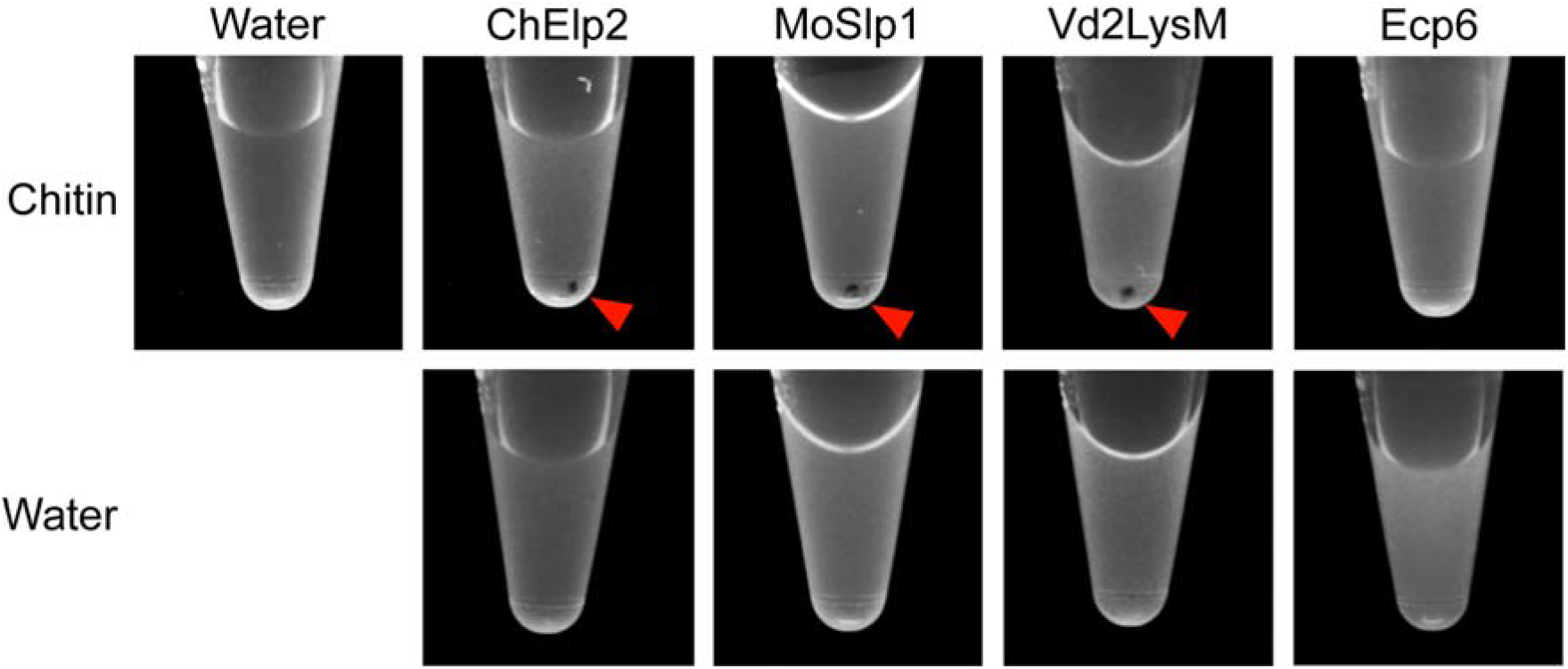
Chitin-induced polymerization of dual-domain LysM effectors. (A) The LysM effector proteins ChElp2, MoSlp1 and Vd2LysM, together with Ecp6 as negative control, were incubated with chitin hexamer or water. After overnight incubation, methylene blue was added and protein solutions were centrifuged, resulting in protein pellets that are indicated by red arrowheads.

## DISCUSSION

We previously determined the mechanism for chitin binding for LysM effectors composed of a single, or of three domains. However, many functionally characterized LysM effectors are composed of two LysMs (Mentlak et al., 2012; Takahara et al., 2016; Kombrink et al., 2017), and it remained unknown whether they are able to form a composite chitin-binding groove through intrachain LysM dimerization to bind chitin with ultrahigh affinity, such as the three-domain LysM effector Ecp6 (Sanchez-Vallet et al., 2013; Sanchez-Vallet et al., 2015). Although our modelling studies based on three software packages quite convincingly eliminated the possibility of intramolecular dimerization, we set out to provide experimental evidence to support this notion and pursued the determination of 3D-protein structures based on X-ray crystallography. We screened the four dual-domain LysM effectors produced in *P. pastoris* as well as in *E. coli* in different concentrations under 500 conditions in absence and presence of chitin in different ratios, as well as with an additive screening in different buffers with a total of nearly 200 conditions, but failed to develop protein crystals. Generally, if crystallization of a protein fails, it can be attributed to many factors, ranging from insufficient purity and homogeneity of the protein, to the fact that some proteins are simply naturally or biologically unable to crystalize (Dessau and Modis, 2011; Wlodawer et al., 2017). However, we then also considered the option that the lack of intramolecular LysM dimerization would perhaps accommodate intermolecular LysM dimerization (Fig. 3A), potentially leading to polymerization, which may lead to polymers of variable lengths and, consequently, compromise protein homogeneity in solution leading to precipitation rather than crystallization in turn. Thus, we decided to investigate the hypothesis that exogenously added chitin induces intermolecular interaction between ChElp2 monomers (Fig. 3B), leading to polymer formation.

Through various lines of experimentation, we now provide solid evidence for the absence of chitin-induced intramolecular dimerization, but rather the occurrence of chitin-induced intermolecular dimerization of LysM domains of ChElp2 and other dual-domain LysM effectors, leading to polymer formation *in vitro*. These experiments include structural predictions, co-immunoprecipitation experiments, DLS measurements of particle size distributions, centrifugation assays, mutant analyses and polymerization assays with various substrates. The finding that Ecp6, in contrast to the dual-domain LysM effectors, does not undergo chitin-induced polymerization is important, as it demonstrates that the pellets are associated with intermolecular dimerization of LysM effector molecules, a process that is not supposed to occur with Ecp6 that undergoes intramolecular LysM dimerization (Sanchez-Vallet et al., 2013). Such protein pellet was also not obtained for the mutant ChElp2^T49W-T120W^ (Fig. 4C), indicating that chitin binding is essential for ChElp2 polymerization. Yet, chitin binding in itself does not lead to protein pellet formation, as limited protein digestion with thermolysin, that cleaves the linker between the two LysMs and releases the individual LysM domains without compromising their chitin-binding ability, did not lead to pellet formation (Fig. 5E). Additionally, the polymerization assays in which ChElp2 was incubated with the chitin dimer, trimer, tetramer, pentamer, hexamer and glucose (Fig. 5) revealed that it is not the mere addition of chitin that leads to precipitation of protein pellets, as only those molecules that have previously been shown to be bound by LysMs resulted in polymerization, further implicating chitin binding as a requirement for the polymerization to occur.

The apparent inability of dual-domain LysM effectors to undergo chitin-induced intramolecular dimerization also implies that these LysM effectors cannot establish ultrahigh affinity chitin-binding sites such as the one that was identified in *C. fulvum* Ecp6 (Sánchez-Vallet et al., 2013). As this finding implies that the chitin binding affinities of the dual-domain LysM effectors is markedly lower than that of Ecp6, it needs to be acknowledged that these LysM effectors may not have the immediate capacity to outcompete host immune receptors for chitin binding. Thus, we speculate that these LysM effectors contribute to fungal virulence *in planta* through the formation of polymeric complexes that have the propensity to precipitate in order to eliminate the presence of freely available chitin oligomers at infection sites that may otherwise bind to the host immune receptors. However, admittedly, experimental evidence for the occurrence of such polymers *in planta* is currently lacking. Previously, we demonstrated that the single LysM domain of Ecp6 that is not involved in the composite ultra-high affinity binding groove, LysM2, can suppress chitin-triggered immunity, indicative of an additional mechanism besides competitive chitin oligosaccharide sequestration (Sánchez-Vallet et al., 2013). Importantly, as argued above, we demonstrated in the current study that Ecp6 does not undergo chitin-induced polymerization. Rather, as proposed previously, LysM2 of Ecp6, but potentially also the dual domain LysM effectors, may perturb the immune receptor dimerization that is required for the activation of immune signaling (<u>Liu et al., 2012</u>), compromising the activation of chitin-triggered immunity in turn (Sánchez-Vallet et al., 2013). To this end, these effectors may physically block host immune receptor dimerization by binding to chitin oligomers that are attached to host immune receptor monomers. Thus, future efforts should be dedicated to discriminate LysM effector precipitation upon polymerization from receptor complex perturbation for the dual-domain LysM effectors when naturally produced by fungi at infection sites during host colonization, momentarily a technically very challenging undertaking.

## MATERIALS AND METHODS

### Sequence alignment and three-dimensional protein structure prediction

LysM domains of proteins were predicted by InterPro (Finn *et al*., 2017) and the alignment of amino acid sequences was performed by ClustalX2. Protein structures were predicted with I-TASSER (Roy et al., 2010; Yang and Zhang, 2015), Phyre2 (Kelley *et al*., 2015) and AlphaFold (Jumper et al., 2021) based on the composite chitin binding groove composed of LysM1 and LysM3 of Ecp6. Structures were viewed by the PyMOL molecular graphics system, version 2 (Schrodinger LLC, 2015).

### Heterologous protein production in *Pichia pastoris*

Protein sequences were analysed using SignalP4.0 (http://www.cbs.dtu.dk/services/SignalP; Petersen *et al*., 2011) and the coding sequences of mature proteins without signal peptide were amplified with primers listed in Table S1, fused with an N-terminal 6×His-tag and cloned into expression vector pPIC9 (Thermo Fisher Scientific, California, USA). Correctness of the resulting constructs was confirmed by DNA sequencing prior to introduction into *Pichia pastoris* strain GS115 (Thermo Fisher Scientific, California, USA). Fermentation was conducted in approximately 3 L of culture in a bioreactor BioFlo120 (Eppendorf, Hamburg, Germany) at 30°C for 5 days, including 3 days of methanol induction. Next, *P. pastoris* cells were pelleted by centrifugation at 3800 g at 4°C for 50 min and the supernatant was concentrated to 200 ml using a Vivaflow 200 Cross Flow Cassette (5000NWCO; Sartorius, Göttingen, Germany) at 4°C for approximately 20 h. The concentrated supernatant was purified using His60 Ni Superflow resin (TaKaRa, California, USA) on a BioLogic LP system (Bio-Rad, California, USA). Purified protein was analysed by protein polyacrylamide gel electrophoresis followed by staining with Coomassie Brilliant Blue (CBB) and dialyzed against 5 L of 50 mM Tris, 150 mM NaCl to remove imidazole. Finally, proteins were further concentrated using Amicon Ultra-15 Centrifugal Filter Units (MERCK, Carrigtohill, Ireland) and stored at -20°C. Final concentrations of the four LysM effectors are listed in Table S2.

### Heterologous protein production in *E. coli*

Coding sequences of mature proteins without signal peptide were amplified with primers listed in Table S1 and cloned into expression vector pETSUMO (Thermo Fisher Scientific, Massachusetts, USA). Correctness of the resulting constructs pETSUMO-ChElp2 and pETSUMO-Vd2LysM were confirmed by DNA sequencing and introduced into *E. coli* strains BL21 and Origami, respectively. FLAG-ChElp2 was amplified using primers FLAG-ChElp2-F and ChElp2-R, cloned into pETSUMO and transformed into BL21. All proteins were produced at 28°C with 0.2 mM IPTG. Cell culture was pelleted by centrifugation at 4000 g for 40 min at 4°C, and the pellet was resuspended in 20 mL lysis buffer (Table S3), shaken at 4°C for at least two hours and centrifuged at 10,000 g for 1 h. The supernatant was collected and purified using His60 Ni Superflow resin (TaKaRa, California, USA) on a BioLogic LP system (Bio-Rad, California, USA). The resulting protein was dialyzed 3 L of 20 mM Tris, 150 mM NaCl, 5 % glycerol, pH 8.0 while 5 µL of cleavage protein ULP1 was added into the dialysis membrane to cleave-off the 6×His-SUMO tag. Next day, protein solution was collected and subjected to purification using His60 Ni Superflow resin (TaKaRa, California, USA) to remove 6×His-SUMO tag from the protein preparations. Eventually, *E. coli*-produced ChElp2 and Vd2LysM were buffer exchanged in 20 mM Tris, 150 mM NaCl, 5% glycerol, pH 8.0 and concentrated to 5 and 4.3 mg/mL respectively.

### Glycoprotein staining assay

1 µL of concentrated *P. pastoris*-produced LysM protein solution was tested using a protein polyacrylamide gel followed by CBB and glycoprotein staining with the Pierce Glycoprotein Staining Kit (Thermo Fisher Scientific, California, USA) according to the manufacturer’s instructions, including the addition of horseradish peroxidase and soybean trypsin inhibitor as positive and negative control, respectively. *E. coli*-produced ChElp2 and Vd2LysM were incubated with chitin hexamer overnight and subjected to glycoprotein staining assay as described above.

### Deglycosylation assay

Deglycosylation was performed PNGase F (MERCK, New Jersey, USA) and Protein Deglycosyaltion Mix II (New England biolabs, Massachusetts, United State) according to the manufacturer’s instructions. Glycosylated fetuin was used as positive control. 5 µL of MoSlp1, Vd2LysM and ChElp2 were treated with 10 µL of PNGase F at 37°C for 16 hours, or with 10 µL of Protein Deglycosylation Mix II at 25°C for 30 min followed by incubation at 37°C for 16 hours. Protein samples were collected at different time points and analysed with protein polyacrylamide gel electrophoresis followed by CBB staining.

### Crystallization conditions

Commercial kits PACT premier™ (Molecular dimensions, Sheffield, UK) and Salt^RX^, Index™, Shotgun, PEG^RX^, PEG/Ion screen (Hampton Research, California, USA) were used for initial screening. 96-well protein crystallization plates were prepared using a Crystal Phoenix robot (Art Robbins Instruments, California, USA). Chitohexaose (Megazyme, Wicklow, Ireland) was added in molar ratios of 3:1 and 1:1. The additive screening was conducted using the Additive Screen HR2-428 (Hampton Research, California, USA) and Tacsimate pH 7.0 (Hampton Research, California, USA) according to the manufacturer’s instructions.

### Co-immunoprecipitation assay

ChElp2 was produced in *E. coli* with either His-sumo-tag or FLAG-tag. 20 µM His-sumo-ChElp2 and 40 µM FLAG-ChElp2 were incubated either individually or together in a total volume of 600 µL in the absence and presence of 40 µM chitohexaose (Megazyme, Wicklow, Ireland) at room temperature for 4-6 h. His-tagged ChElp2 was pulled-down with Dynabeads™ His-Tag (Thermo Fisher Scientific, Massachusetts, USA) and the subsequent western blot analyses were performed with anti-His (Sigma, Missouri, USA) or anti-FLAG antibody (Sigma, Missouri, USA).

### Isothermal titration calorimetry

Isothermal titration calorimetry (ITC) experiments were performed at 25°C using a MicroCal PEAQ-ITC Automated (Malvern Panalytical, Worcestershire, UK) according to the manufacturer’s instructions. *E. coli*-produced LysM proteins were buffer exchanged in 50 mM Tris, 150 mM NaCl, 5% glycerol, pH 8.0. ChElp2 (20 µM) and ChElp2^T49W-T120W^ (34 µM) were titrated with a first injection of 0.5 µL chitohexaose (Megazyme, Wicklow, Ireland) with a concentration of 200 µM that was prepared in the same buffer with LysM proteins, followed by another 20 injections of 2 µL. Data were analysed using Origin (OriginLab, Northampton, MA, USA) and fitted to a one-binding-site model.

### Dynamic light scattering (DLS) measurements

LysM proteins were dialyzed against 20 mM Tris, 150 mM NaCl, pH 8.0 overnight and treated with 0.1% Triton X-100. Chitohexaose (Megazyme, Wicklow, Ireland) was incubated with LysM proteins in protein:chitin molar ratios of 1:0, 1:3 and 1:6 overnight. Next day, particle size distribution was measured with a Zetasizer Nano (Malvern Panalytical, Worcestershire, UK). The data was visualized with Zetasizer software.

### Polymerization assays

LysM effector proteins were adjusted to a concentration of 200 µM and 200 µL of each protein was incubated with 200 µL of 2 mM diacetyl-chitobiose (Chi2), triacetyl-chitobiose (Chi3), tetraacetyl-chitotetraose (Chi4), pentaacetyl-chitopentaose (Chi5), hexaacetyl-chitohexaose (Chi6) (Megazyme, Wicklow, Ireland), glucose (Merck, Darmstadt, Germany), or 200 µL water as control, at room temperature overnight. The next day, 2 µL of 0.2% methylene blue (Sigma-Aldrich, Missouri, USA) was added and incubated for 30 min after which protein solutions were centrifuged at 20,000 g for 15 min. Photos were taken with a ChemiDoc XRS+ system (Bio-Rad, California, USA) with custom setting for RFP or with a ChemiDoc system (Bio-Rad, California, USA) with the setting of Coomassie Blue with white tray (Bio-rad, California, USA).

For limited digestion assay with thermolysin, 10 µL of ChElp2 (200 µM) was incubated with 5 µL of 1 mg/ml thermolysin at 55°C for 3 hours. Protein samples were collected at different time points and analysed with a protein polyacrylamide gel. Subsequently, ChElp2 was digested with thermolysin for 1 h, deactivated the protease according to manufacturer’s instructions and incubated with chitin hexamer overnight. The next day, 2 µL of 0.2% methylene blue (Sigma-Aldrich, Missouri, USA) was added and incubated for 30 min. After centrifugation at 20,000 g for 15 min, photos were taken with a ChemiDoc system (Bio-Rad, California, USA) with the setting of Coomassie Blue with white tray (Bio-rad, California, USA).

## ACKNOWLEDGEMENTS

H.T. acknowledges receipt of a PhD fellowship from the China Scholarship Council (CSC) and G.L.F acknowledges receipt of a PhD fellowship from the Coordination for the Improvement of Higher Education Personnel (CAPES) from the federal government of Brazil. B.P.H.J.T acknowledges funding by the Alexander von Humboldt Foundation in the framework of an Alexander von Humboldt Professorship endowed by the German Federal Ministry of Education and Research, and is furthermore supported by the Deutsche Forschungsgemeinschaft (DFG, German Research Foundation) under Germany’s Excellence Strategy – EXC 2048/1 – Project ID: 390686111. The authors declare no conflict of interest exists.

## AUTHOR CONTRIBUTIONS

HT, JRM, BPHJT conceived the study; HT, JRM, BPHJT designed experiments; HT, GLF and AK performed experiments; HT, JRM, BPHJT analyzed data and wrote the manuscript; JRM and BPHJT supervised the project; all authors discussed the results and contributed to the final manuscript.

## REFERENCES

1. Altenbach D, Robatzek S (2007) Pattern recognition receptors: from the cell surface to intracellular dynamics. Mol Plant-Microbe Interact 20: 1031–1039

2. Baker HM, Day CL, Norris GE, Baker EN (1994) Enzymatic deglycosylation as a tool for crystallization of mammalian binding proteins. Acta Crystallogr D Biol Crystallogr 50: 380–384

3. Bateman A, Bycroft M (2000) The structure of a LysM domain from E. coli membrane-bound lytic murein transglycosylase D (MltD). J Mol Biol 299: 1113–1119

4. Boller T, Felix G (2009) A renaissance of elicitors: perception of microbe-associated molecular patterns and danger signals by pattern-recognition receptors. Annu Rev Plant Biol 60: 379–406

5. Bolton MD, Van Esse HP, Vossen JH, De Jonge R, Stergiopoulos I, Stulemeijer IJE, Van Den Berg GCM, Borrás-Hidalgo O, Dekker HL, De Koster CG, et al (2008) The novel Cladosporium fulvum lysin motif effector Ecp6 is a virulence factor with orthologues in other fungal species. Mol Microbiol 69: 119–136

6. Cao Y, Liang Y, Tanaka K, Nguyen CT, Jedrzejczak RP, Joachimiak A, Stacey G (2014) The kinase LYK5 is a major chitin receptor in Arabidopsis and forms a chitin-induced complex with related kinase CERK1. Elife 3: e03766

7. Cen K, Li B, Lu Y, Zhang S, Wang C (2017) Divergent LysM effectors contribute to the virulence of Beauveria bassiana by evasion of insect immune defenses. PLOS Pathog 13: e1006604

8. Chen XL, Shi T, Yang J, Shi W, Gao X, Chen D, Xu X, Xu JR, Talbot NJ, Peng YL (2014) N-Glycosylation of effector proteins by an α-1,3-Mannosyltransferase is required for the rice blast fungus to evade host innate immunity. Plant Cell 26: 1360–1376

9. Davis SJ, Puklavec MJ, Ashford DA, Harlos K, Jones EY, Stuart DI, Williams AF (1993) Expression of soluble recombinant glycoproteins with predefined glycosylation: application to the crystallization of the T-cell glycoprotein CD2. Protein Eng 6: 229–232

10. Dessau MA, Modis Y (2011) Protein crystallization for X-ray crystallography. J Vis Exp 16: 2285

11. Dolfors F, Holmquist L, Dixelius C, Tzelepis G (2019) A LysM effector protein from the basidiomycete Rhizoctonia solani contributes to virulence through suppression of chitin-triggered immunity. Mol Genet Genomics 294: 1211–1218

12. Felix G, Regenass M, Boller T (1993) Specific perception of subnanomolar concentrations of chitin fragments by tomato cells: Induction of extracellular alkalinization, changes in protein phosphorylation, and establishment of a refractory state. Plant J 4: 307–316

13. Finn RD, Attwood TK, Babbitt PC, Bateman A, Bork P, Bridge AJ, Chang HY, Dosztanyi Z, El-Gebali S, Fraser M, et al (2017) InterPro in 2017-beyond protein family and domain annotations. Nucleic Acids Res 45: D190–D199

14. Free SJ (2013) Fungal cell wall organization and biosynthesis. Adv Genet 81: 33–82

15. Haltiwanger RS, Lowe JB (2004) Role of glycosylation in development. Annu Rev Biochem 73: 491–537

16. Jones JDG, Dangl JL (2006) The plant immune system. Nature 444: 323–329

17. de Jonge R, van Esse P, Kombrink A, Shinya T, Desaki Y, Bours R, van der Krol S, Shibuya N, Joosten MHAJ, Thomma BPHJ (2010) Conserved fungal LysM effector Ecp6 prevents chitin-triggered immunity in plants. Science (80-) 329: 953–955

18. Jumper J, Evans R, Pritzel A, Green T, Figurnov M, Ronneberger O, Tunyasuvunakool K, Bates R, Žídek A, Potapenko A, et al (2021) Highly accurate protein structure prediction with AlphaFold. Nature 596: 583–589

19. Kelley LA, Mezulis S, Yates CM, Wass MN, Sternberg MJE (2015) The Phyre2 web portal for protein modeling, prediction and analysis. Nat Protoc 10: 845–858

20. Kombrink A, Rovenich H, Shi-Kunne X, Rojas-Padilla E, van den Berg GCM, Domazakis E, de Jonge R, Valkenburg DJ, Sánchez-Vallet A, Seidl MF, et al (2017) Verticillium dahliae LysM effectors differentially contribute to virulence on plant hosts. Mol Plant Pathol 18: 596–608

21. Kombrink A, Thomma BPHJ (2013) LysM effectors: secreted proteins supporting fungal life. PLOS Pathog 9: e1003769

22. Lenardon MD, Munro CA, Gow NA (2010) Chitin synthesis and fungal pathogenesis. Curr Opin Microbiol 13: 416–423

23. Liu T, Liu Z, Song C, Hu Y, Han Z, She J, Fan F, Wang J, Jin C, Chang J, Zhou JM, Chai J. (2012) Chitin-induced dimerization activates a plant immune receptor. Science 336: 1160–1164

24. Marshall R, Kombrink A, Motteram J, Loza-Reyes E, Lucas J, Hammond-Kosack KE, Thomma BPHJ, Rudd JJ (2011) Analysis of two in planta expressed LysM effector homologs from the fungus Mycosphaerella graminicola reveals novel functional properties and varying contributions to virulence on wheat. Plant Physiol 156: 756–769

25. Mentlak TA, Kombrink A, Shinya T, Ryder LS, Otomo I, Saitoh H, Terauchi R, Nishizawa Y, Shibuya N, Thomma BPHJ, et al (2012) Effector-mediated suppression of chitin-triggered immunity by Magnaporthe oryzae is necessary for rice blast disease. Plant Cell 24: 322–335

26. Miya A, Albert P, Shinya T, Desaki Y, Ichimura K, Shirasu K, Narusaka Y, Kawakami N, Kaku H, Shibuya N (2007) CERK1, a LysM receptor kinase, is essential for chitin elicitor signaling in Arabidopsis. Proc Natl Acad Sci U S A 104: 19613–19618

27. Moremen KW, Tiemeyer M, Nairn A V (2012) Vertebrate protein glycosylation: diversity, synthesis and function. Nat Rev Mol Cell Biol 13: 448–462

28. Nagashima Y, von Schaewen A, Koiwa H (2018) Function of N-glycosylation in plants. Plant Sci 274: 70–79

29. Petersen TN, Brunak S, von Heijne G, Nielsen H (2011) SignalP 4.0: discriminating signal peptides from transmembrane regions. Nat Methods 8: 785–786

30. Petutschnig EK, Jones AME, Serazetdinova L, Lipka U, Lipka V (2010) The Lysin Motif Receptor-like Kinase (LysM-RLK) CERK1 is a major chitin-binding protein in Arabidopsis thaliana and subject to chitin-induced phosphorylation. J Biol Chem 285: 28902–28911

31. Proteau A, Shi R, Cygler M (2010) Application of dynamic light scattering in protein crystallization. Curr Protoc Protein Sci Chapter 17: Unit 17 10

32. Rovenich H, Boshoven JC, Thomma BPHJ (2014) Filamentous pathogen effector functions: Of pathogens, hosts and microbiomes. Curr Opin Plant Biol 20: 96–103

33. Rovenich H, Zuccaro A, Thomma BP (2016) Convergent evolution of filamentous microbes towards evasion of glycan-triggered immunity. New Phytol 212: 896–901

34. Roy A, Kucukural A, Zhang Y (2010) I-TASSER: a unified platform for automated protein structure and function prediction. Nat Protoc 5: 725–738

35. Sanchez-Vallet A, Mesters JR, Thomma BPHJ (2015) The battle for chitin recognition in plant-microbe interactions. FEMS Microbiol Rev 39: 171–183

36. Sanchez-Vallet A, Saleem-Batcha R, Kombrink A, Hansen G, Valkenburg DJ, Thomma BPHJ, Mesters JR (2013) Fungal effector Ecp6 outcompetes host immune receptor for chitin binding through intrachain LysM dimerization. Elife 2: e00790

37. Sánchez-Vallet A, Tian H, Rodriguez-Moreno L, Valkenburg D-J, Saleem-Batcha R, Wawra S, Kombrink A, Verhage L, de Jonge R, van Esse HP, et al (2020) A secreted LysM effector protects fungal hyphae through chitin-dependent homodimer polymerization. PLoS Pathog 16: 1–21

38. Schrodinger LLC (2015) The JyMOL Molecular Graphics Development Component, Version 1.8.

39. Takahara H, Hacquard S, Kombrink A, Hughes HB, Halder V, Robin GP, Hiruma K, Neumann U, Shinya T, Kombrink E, et al (2016) Colletotrichum higginsianum extracellular LysM proteins play dual roles in appressorial function and suppression of chitin-triggered plant immunity. New Phytol 211: 1323–1337

40. Tang J, Sun Y, Han Z, Shi W (2019) An illustration of optimal selected glycosidase for N-glycoproteins deglycosylation and crystallization. Int J Biol Macromol 122: 265–271

41. Tian H, MacKenzie CI, Rodriguez-Moreno L, van den Berg GCM, Chen H, Rudd JJ, Mesters JR, Thomma BPHJ (2021) Three LysM effectors of Zymoseptoria tritici collectively disarm chitin-triggered plant immunity. Mol Plant Pathol 1–11

42. Wlodawer A, Dauter Z, Jaskolski M (2017) Protein crystallography: methods and protocols. Humana Press 1607: 672

43. Xu C, Ng DT (2015) Glycosylation-directed quality control of protein folding. Nat Rev Mol Cell Biol 16: 742–752

44. Yang J, Zhang Y (2015) I-TASSER server: new development for protein structure and function predictions. Nucleic Acids Res 43: W174–81

45. Zeng T, Rodriguez-Moreno L, Mansurkhodzaev A, Wang P, van den Berg W, Gasciolli V, Cottaz S, Fort S, Thomma B, Bono JJ, et al (2020) A lysin motif effector subverts chitin-triggered immunity to facilitate arbuscular mycorrhizal symbiosis. New Phytol 225: 448–460

46. Zhang X-C, Wu X, Findley S, Wan J, Libault M, Nguyen HT, Cannon SB, Stacey G (2007) Molecular Evolution of Lysin Motif-Type Receptor-Like Kinases in Plants. Plant Physiol 144: 623–636

47. Zipfel C (2008) Pattern-recognition receptors in plant innate immunity. Curr Opin Immunol 20: 10–16

