## Supplemental Information for "Fungal dual-domain LysM effectors undergo chitin-induced intermolecular, and not intramolecular, dimerization"

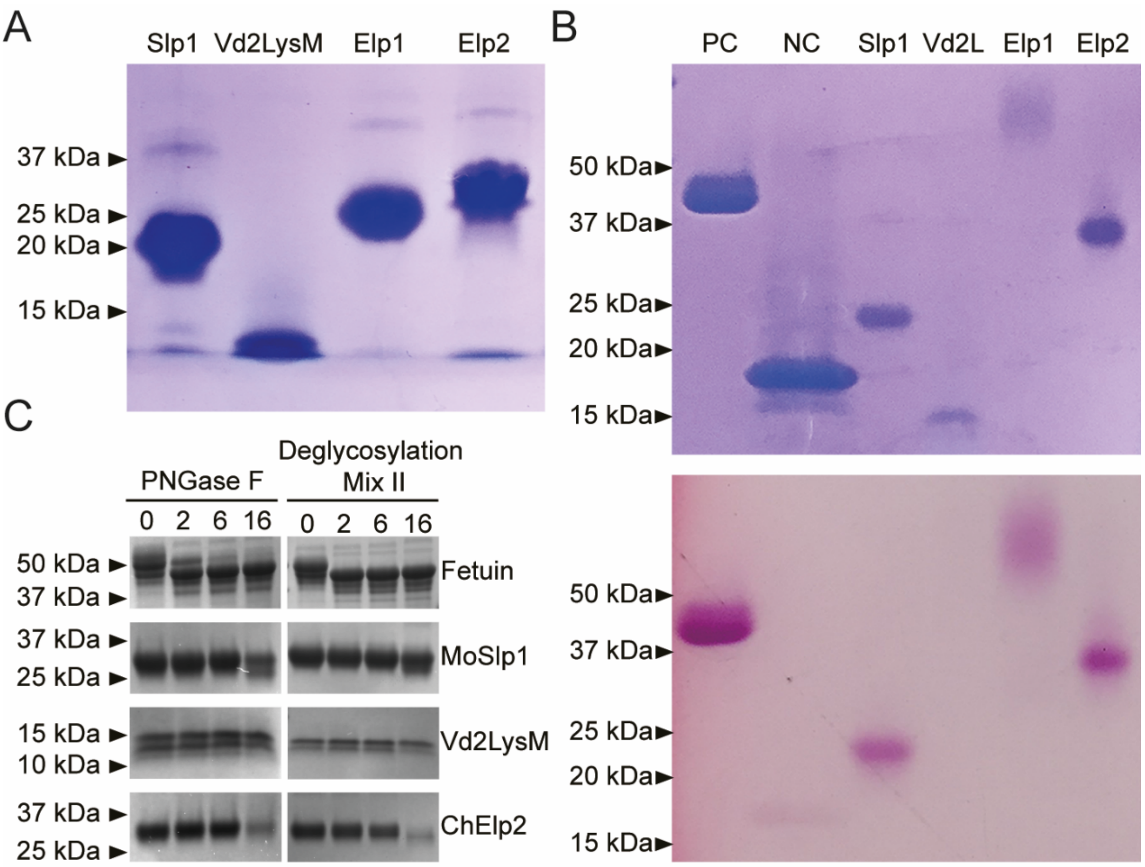


**Fig. S1 Heterologous production of the four *P. pastoris*-produced dual-domain LysM effectors.** (A) Protein polyacrylamide gel electrophoresis of 1 µL of the purified *Pichia pastoris*-produced dual LysM domain effectors *Magnaporthe oryzae* MoSlp1, *Verticillium dahliae* Vd2LysM, and *Colletotrichum higginsianum* ChElp1 and ChElp2 followed by Coomassie brilliant blue (CBB) staining. (B) CBB (upper panel) and glycoprotein staining (bottom panel) of polyacrylamide gels with *Pichia pastoris*-produced dual LysM domain effectors. PC and NC are positive (horseradish peroxidase) and negative (soybean trypsin inhibitor) control, respectively. (C) Deglycosylation assay performed with peptide:N-glycosidase F (PNGase F) and Protein Deglycosylation Mix II on MoSlp1, Vd2LysM, ChElp2 or fetuin as positive control. Protein samples were collected after different incubation times and subjected to polyacrylamide gel electrophoresis followed by CBB staining.

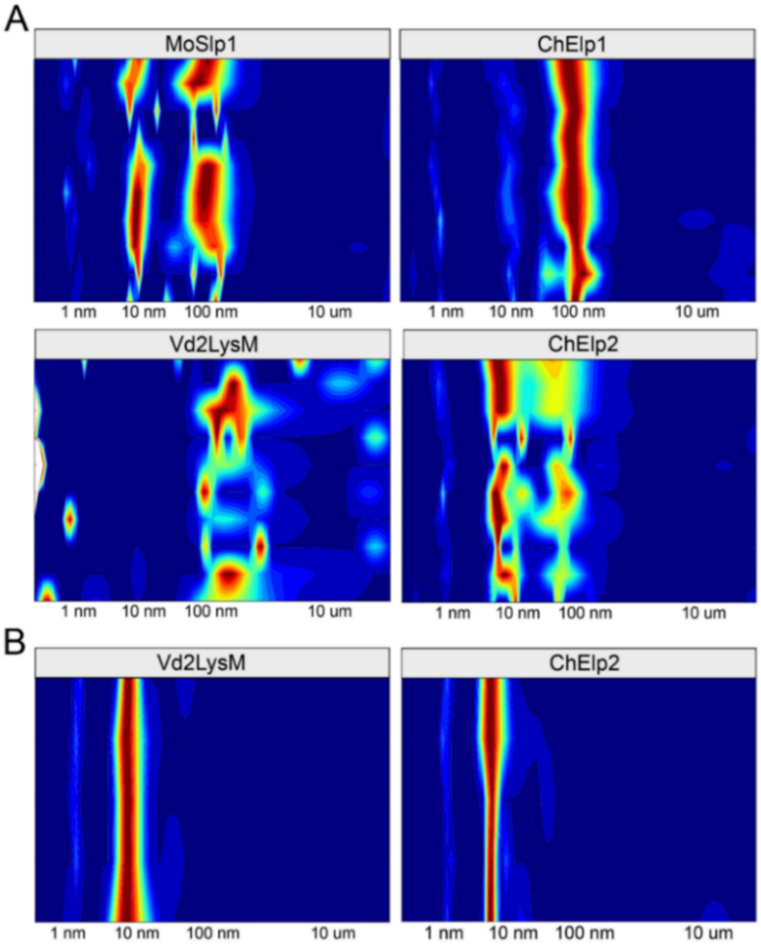


**Fig. S2 Particle size distribution of dual-domain LysM effectors as determined by dynamic light scattering (DLS).** The particle size distribution is shown as a colour scale heat map ranging from blue (lowest abundance) to red (highest abundance) for a particle size range of 1 nm to 100 µM. (A) Heat maps of the four *Pichia pastoris*-produced LysM effectors after initial purification and concentration. (B) Heat maps of Vd2LysM and ChElp2 after gel filtration and decyl β-D-maltopyranoside (DM) treatment.

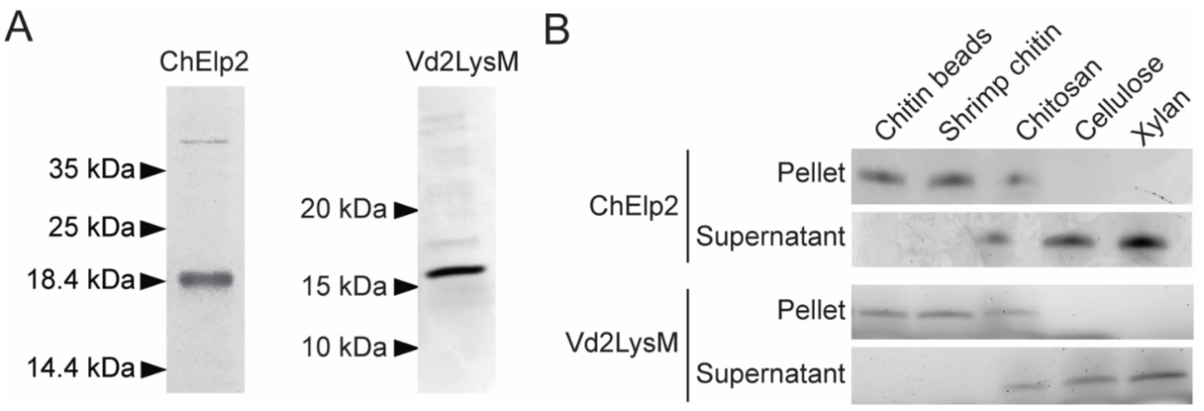


**Fig. S3 *E. coli*-produced ChElp2 and Vd2LysM bind chitin.** (A) ChElp2 and Vd2LysM were produced in *E. coli* and their quality was evaluated on protein polyacrylamide gel. (B) *E. coli*-produced ChElp2 or Vd2LysM were incubated with insoluble magnetic chitin beads, shrimp shell chitin, chitosan, cellulose and xylan for 6 hours. After centrifugation, pellet and supernatant fractions were analyzed on a protein polyacrylamide gel.

**Table S1. Primers used in this study.**

| Primers | Sequences |
|---|---|
| His-Flag-MoSlp1-F | CGGTATGAATTC**ATG**CATCATCATCATCATCAT CCCGACTACAAGGACGACGATGACAAGGCCATGCCTCAGGCAAC |
| His-Flag-MoSlp1-R | CGGTATGCGGCCGCCTAGTTCTTGCAGATGGGGATGTTG |
| His-Flag-Vd2L-F | CGGTATGAATTC**ATG**CATCATCATCATCATCAT CCCGACTACAAGGACGACGATGACAAGTACCGAAGGACATGCAGTCATACC |
| His-Flag-Vd2L-R | CGGTATGCGGCCGCTTAGTTCCAGCTGCACGGC |
| His-Flag-ChElp1-F | CGGTATGAATTC**ATG**CATCATCATCATCATCAT CCCGACTACAAGGACGACGATGACAAGCTCCCTCAGGCTACCCCGACCACG |
| His-Flag-ChElp1-R | CGGTATGCGGCCGCTTAACAGACGGGGATGTTGATGACATCGC |
| His-Flag-ChElp2-F | CGGTATGAATTC**ATG**CATCATCATCATCATCAT CCCGACTACAAGGACGACGATGACAAGCTCCCTGTCTGTGACGCCACTAC |
| His-Flag-ChElp2-R | CGGTATGCGGCCGCCTAACACACAGGGACGTTGATGACC |
| MoSlp1-F | CGGTATGAATTCGCCATGCCTCAGGCAAC |
| MoSlp1-R | CGGTATGCGGCCGC**TTATTA**CTAGTTCTTGCAGATGGGGATGTTG |
| Vd2L-F | CGGTATGAATTCTACCGAAGGACATGCAGTCATACC |
| Vd2L-R | CGGTATGCGGCCGC**TTATTA**TTAGTTCCAGCTGCACGGC |
| ChElp1-F | CGGTATGAATTCCTCCCTCAGGCTACCCCGACCACG |
| ChElp1-R | CGGTATGCGGCCGC**TTATTA**TTAACAGACGGGGATGTTGATGACATCGC |
| ChElp2-F | CGGTATGAATTCCTCCCTGTCTGTGACGCCACTAC |
| ChElp2-R | CGGTATGCGGCCGC**TTATTA**CTAACACACAGGGACGTTGATGACC |
| FLAG-ChElp2-F | CGGTATGAATTCCCCGACTACAAGGACGACGATGACAAGCTCCCTGTCTGTGACGCC ACTAC |

*The coding sequence for the 6×His-tag is indicated with cyan font, the coding sequence for the FLAG-tag is indicated with green font, and restriction enzyme recognition sites are indicated with red font.

**Table S2. Final concentrations of the four *P. pastoris*-produced LysM effectors.**

| Protein | Concentration |
|---|---|
| MoSlp1 | 10 mg/ml |
| Vd2LysM | 7.35 mg/ml |
| ChElp1 | 9.0 mg/ml |
| ChElp2 | 14 mg/ml |

**Table S3. Lysis buffer composition (20 mL).**

| Amount | Reagent | Company |
|---|---|---|
| 18 mL | 50 mM Tris, 150 nM NaCl, pH 8.5 | |
| 2 mL | Glycerol | VWR, Ohio, USA |
| 120 mg | Lysozyme from chicken egg white | Sigma-Aldrich, Missouri, USA |
| 80 mg | Sodium deoxycholate | Sigma-Aldrich, Missouri, USA |
| 1.25 mg | Deoxyribonuclease I | Sigma-Aldrich, Missouri, USA |
| 1 pill | Protease inhibitor cocktail | Roche, Mannheim, Germany |
